# Cockroaches show individuality in learning and memory during classical and operant conditioning

**DOI:** 10.1101/825265

**Authors:** Cansu Arican, Janice Bulk, Nina Deisig, Martin Paul Nawrot

## Abstract

Animal personality and individuality are intensively researched in vertebrates and both concepts are increasingly applied to behavioral science in insects. However, only few studies have looked into individuality with respect to performance in learning and memory tasks. In vertebrates individual learning capabilities vary considerably with respect to learning speed and learning rate. Likewise, honeybees express individual learning abilities in a wide range of classical conditioning protocols. Here, we study individuality in the learning and memory performance of cockroaches, both in classical and operant conditioning tasks. We implemented a novel classical (olfactory) conditioning paradigm where the conditioned response is established in the maxilla-labia response (MLR). Operant spatial learning was investigated in a forced two-choice task using a T-maze. Our results confirm individual learning abilities in classical conditioning of cockroaches that was reported for honeybees and vertebrates but contrast long-standing reports on stochastic learning behavior in fruit flies. In our experiments, most learners expressed a correct behavior after only a single learning trial showing a consistent high performance during training and test. We can further show that individual learning differences in insects are not limited to classical conditioning but equally appear in operant conditioning of the cockroach.

## Introduction

A behavioral syndrome defines a consistent behavior of an individual that is correlated across time and contexts. Animal personality (Gosling and Vazire, 2002) is expressed in long-term differences among individuals across a variety of behavioral traits such as boldness– shyness, exploration–avoidance, activity level, sociability or aggression (Dingemanse and Wolf, 2010; Sih et al., 2004b, 2004a). While consistent behavioral traits have been heavily studied in vertebrates, literature on individuality and personality in invertebrates is still scarce (for review see Kralj-Fišer and Schuett, 2014). The small amount of available data on invertebrate personality may be partly due to the traditional belief that invertebrates express stereotyped stimulus-response behaviors with little individual differences (e.g. Brembs, 2013). Invertebrate studies have primarily been conducted in the context of collective behavior in social contexts and mostly investigated how individual personalities influence the colony behavior (e.g. in cockroaches: Planas-Sitjà et al., 2018; Planas-Sitjà and Deneubourg, 2018, in ants: Pinter-Wollman, 2012, spiders: Grinsted et al., 2013; Wright et al., 2014, pea aphids: Schuett et al., 2011; and crickets: Rose et al., 2017).

At the level of animal cognition, inter-individual performance differences may reflect variation in cognitive ability independent of animal personality. However, individual cognitive styles may also inflict personality (Carere and Locurto, 2011). Individuality has been intensively studied in learning and memory. Learning and memorizing the relevance of environmental cues is of major importance for the survival of virtually all animals. Individuals of a species can vary substantially in their learning performances as has been shown for both vertebrates (e.g. David et al., 2011; Gallistel et al., 2004; Gosling, 2001; Groothuis and Carere, 2005; Kolata et al., 2005; Kotrschal and Taborsky, 2010; Schuett and Dall, 2009) and invertebrates (for review see Dukas, 2008).

In insects, studies have focused on bumble bees and honey bees. Bumble bees have been studied in a variety of tasks (Chittka et al., 2003). For example, individual bumble bees, which learn only a single flower parameter (odor or color) were more efficient in several ways than those, which had learned two: they made fewer errors, had shorter flower handling times, corrected errors faster, and transitions between flowers were initially more rapid (Chittka and Thomson, 1997). It has further been shown that individual bumblebees consistently differ in their ability to learn to discriminate stimuli from the visual and olfactory modality (Muller and Chittka, 2012). A systematic analysis of classical conditioning experiments in the honeybee found that the group-average learning behavior did not adequately represent the behavior of individual animals. This result was consistent across a large number of datasets including olfactory and tactile conditioning collected from more than 3000 honeybees obtained during absolute and differential classical conditioning (Pamir et al., 2011, 2014). Gradually increasing learning curves reflected an artifact of group averaging and the behavioral performance of individuals was characterized by an abrupt and often step-like increase in the level of response (Pamir et al., 2011), a result that directly matches observation in vertebrates (Gallistel et al., 2004) but contradicts earlier findings in the fruit fly (*Drosophila melanogaster*) in which the group-average behavior has been described to represent the probabilistic expression of behavior in individuals (Quinn et al., 1974).

In the present work, we asked whether cockroaches show individuality in their learning performances, both in classical and operant conditioning tasks. Behavioral learning studies that used olfactory and visual cues demonstrated that cockroaches can be assayed for classical conditioning tasks while animals are immobilized (Kwon et al., 2004; Lent and Kwon, 2004; Watanabe et al., 2003; Watanabe and Mizunami, 2006) or able to move freely (Hosono et al., 2016; Sato et al., 2006; Watanabe et al., 2003). In some experiments, after classical olfactory conditioning memory tests were performed in an open arena where cockroaches could freely choose to approach different odors (Sato et al., 2006; Watanabe et al., 2003). Open arenas and T-mazes have been used successfully for operant conditioning in cockroaches. Balderrama (1980) demonstrated for the first time that cockroaches could be trained to associate different odors with either sugar or salt solution in an open arena. Mizunami et al (1998) studied place memory using a spatial heat maze with and without visual cues. More recent studies by Mizunami and colleagues (Sakura et al., 2002; Sakura and Mizunami, 2001) confirmed and extended operant conditioning of cockroaches in the open arena. Barraco et al (1981) successfully trained cockroaches in a spatial discrimination task using an electric shock to punish either a left or right turn in a T-maze. Employing stimuli of different modalities, we show in the present study that cockroaches demonstrate individuality in their ability to learn and memorize stimuli employing both classical and operant conditioning task.

## Material & Methods

### Insects

All experiments were conducted with adult male *Periplaneta americana*. Animals were kept under a reverse light-dark cycle (12h:12h) at 26°C in laboratory colonies at our rearing facilities at the University of Cologne. Cockroaches were allowed to drink water and fed on oat *ad libitum*. However, water was removed four days before training to increase motivation. All experiments were conducted in the active phase (scotophase) of the animals.

### Experimental setups

For classical conditioning cockroaches were harnessed in custom-made fixation cylinders (Figure 1A) after anaesthetization at 4°C. After fixation, only the animals’ head protruded allowing free movement of the antennae and mouthparts. After habituation in the experimental room for 16 h, cockroaches were placed in front of a 10 ml plastic syringe mounted in a holder and an exhaust system behind removing odor-loaded air (Figure 1A). Odors were diluted in mineral oil (Acros Organics^™^, Geel, Belgium) and odor concentrations were adjusted to match the vapor pressure of the odor with the lowest value (trans-cinnamaldehyde). Dilutions were as follows (in % v/v): isoamyl acetate (99+ %, pure, Acros Organics^™^, Geel, Belgium): 26.27 %, butyric acid (> 99 %, Aldrich, Darmstadt, Germany): 2.56 %, as well trans-cinnamaldehyde (≥ 98 %, Merck, Darmstadt, Germany) undiluted. Ten µl of each odor were given on a piece of filter paper inserted in a 10 ml plastic syringe for olfactory stimulation. A filter paper without any odor nor the solvent was used as control stimulus testing for mechanical stimulation (Air). Isoamyl acetate and butyric acid were used as conditioned stimuli (CS+ or CS-), while trans-cinnamaldehyde served as control odor without any assigned contingency (reward or punishment). Odors were chosen based on choice behavior of cockroaches in preliminary tests in a T-maze, in which no preference was found between isoamyl acetate and butyric acid.

**Figure 1.**
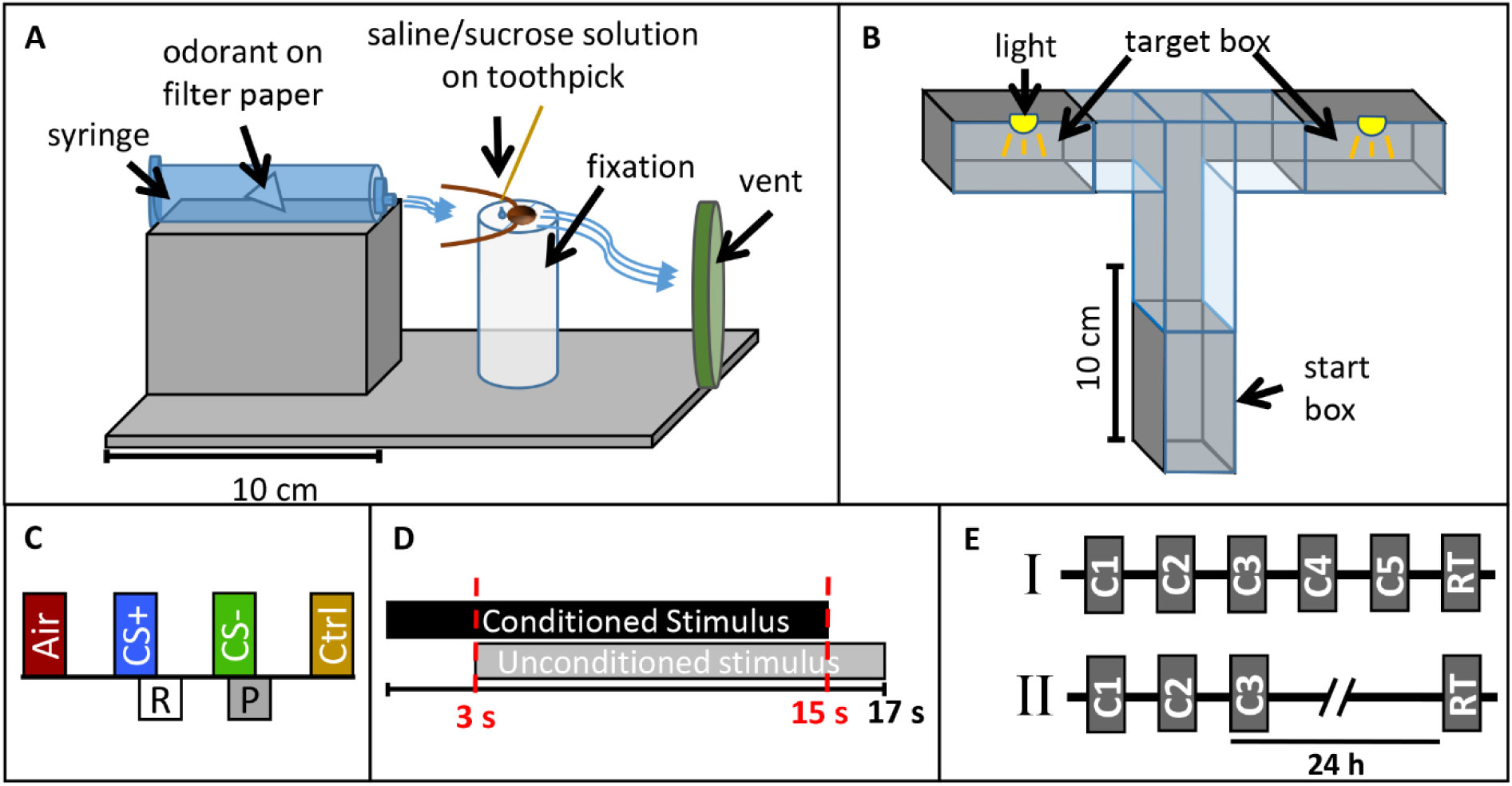
Experimental design. Sketch of A) the classical conditioning setup and B) the operant conditioning setup. Presentation of one training block in the classical conditioning paradigm. Each block contains an air puff (Air), the rewarded odor CS+ paired with sugar solution (R), the punished odor CS- paired with saline solution (P) and the control odor (Ctrl, cinnamaldehyde). D) Time sequence of the conditioned stimulus (CS, odor) and unconditioned stimulus (US, sucrose or saline solution) in a single training trial for classical conditioning. E) Operant conditioning protocols I) with five consecutive training trials (C1-C5) and a retention test (RT) after 35 min and II) with three consecutive training trials (C1-C3) and a retention test after 24 h.

For operant conditioning, a custom-made flexible maze was used. Walls made from polyvinyl chloride allowed easy cleaning with alcohol between single trials. The maze was positioned on a ground plate and different tunnel pieces were combined to form a T-Maze (Figure 1B). Shutters allowed closing the start and target boxes (20 cm × 28 cm × 4 cm). All experiments with the T-maze were conducted under red light (Figure 1B).

### Training and test procedures

First, we established a novel classical conditioning paradigm in the harnessed cockroach, training the animals to exhibit a specific movement of the maxilla-labia (mouthparts) as conditioned response behavior. We termed this response the maxilla-labia response (MLR). When touching the antennae and mouthparts with sucrose solution, cockroaches start to quickly move and extend their maxillae and labium, the most central mouthparts, to reach for and suck the solution. When saline solution is presented, animals touch and taste the solution without ingesting and show clear avoidance behavior (retraction of the mouthparts). In each single trial the occurrence or non-occurrence of the MLR was recorded as a binary response (0/1). Only if the MLR was observed within the first three seconds of odor presentation (before US-onset, see Figure 1C and description below) it was counted as conditioned response. This novel paradigm for classical conditioning of the cockroach is similar to the proboscis extension response paradigm used in classical conditioning of honeybees, first established by Takeda (1961) and later standardized by Bitterman et al. (1983).

For classical conditioning, each block of training consisted of i) one stimulation with a simple filter paper without an odorant to test for a mechanical response to the air puff (Air), ii) one CS+ presentation (reinforced conditioned stimulus) paired with 20 % sucrose solution as positive reinforce (unconditioned stimulus, US), iii) one CS-presentation (punished stimulus) paired with 20 % saline solution as negative reinforce, as well as iv) one stimulation with a control odor (cinnamaldehyde, Ctrl), which was not associated to a US (Figure 1C). In each CS+ or CS-presentation the respective odor (CS) was presented for 15 s. Three seconds after odor onset, the US was delivered by touching the maxillary palps with sucrose or saline solution and animals were allowed to drink the respective solutions for 14 s (Figure 1D). In the case of the negative reinforcer, most animals did not drink the saline solution voluntarily but were ‘forced’ to taste the salt in all trials by touching their mouthparts with the toothpick. We performed all experiments in two independent groups for which the identities of the CS+ odor and the CS- was reversed. For retention tests the same pattern of odor presentation as in conditioning trials was used except that no US were presented.

Three differential classical conditioning experiments were conducted. In each block of training trials, the two control odors (air, cinnamaldehyde) were separated from the two CSs with an interstimulus interval (ISI) of 45 s, whereas the ISI between CS+ and CS- was 32 s. The first experiment was designed to investigate differential learning with an acquisition phase that consisted of five blocks of trials (each block contained one presentation of air, the CS+, the CS- and a control stimulation, respectively) with an inter-block interval of 10 min. A retention test was conducted after 10 min. The second and third experiments were designed to investigate memory retention after differential learning at two different time intervals (1 h and 24 h). Due to the length of the experiment, only three training blocks with an inter-block interval of 10 min were used for these two experiments.

For operant conditioning, each cockroach was allowed to acclimate in the start box for 15 min before training. At the beginning of a training trial, the shutter was opened and the cockroaches were allowed to walk freely and enter the target boxes. When entering one of the target boxes for the first time, the shutter was closed and the animal was subjected to a 5 min light exposure (punished stimulus, US). Whenever an animal entered the same target box again in a subsequent trial, it was again subjected to the light punishment. All animals which did not start moving within the first 3 min in two consecutive attempts were excluded from the experiment.

Two different operant conditioning paradigms were used. In the first one, animals were trained in five blocks of trials with an inter trial interval (ITI) of 35 min and memory retention was tested after 35 min. In the second, animals were trained in three blocks of trials with an ITI of 35 min and a retention test was performed 24 h later (Figure 1E).

### Statistics

The results were analyzed with Matlab R2019a (The MathWorks, Natick, Massachusetts, USA) and IBM SPSS Statistics Version 25.0 (IBM Corp., Armonk, New York, USA) and visualized with Matlab R2019a and Photoshop (Adobe Inc., San José, California, USA).

We analyzed spontaneous responses to different odors in two groups of animals. We pooled the behavioral response to odor presentations in the first training trial and before US presentation across all individuals that had been treated in parallel and under identical experimental conditions. Chi^2^ tests were used to compare responses to different odors. Additionally, we calculated the Phi coefficient to analyze the correlation between odor responses across individuals.

We analyzed the classical conditioning experiments by dividing them according to odors that were used as CS+ and CS-. In addition, we pooled all animals with the same conditioning pattern regardless of the odor that was used as CS+ or CS-. To analyze the response to CS+ we excluded all animals showing spontaneous responses to the CS+ in the first trial. For analyzing responses to the CS- we excluded all animals that did not respond to it in the first trial. This is a common procedure to exclude spontaneously responding animals and to visualize the learning curve (Giurfa and Sandoz, 2012; Pamir et al., 2011). For the statistical analysis of the classical conditioning experiments we applied three different statistical tests. First, one-way ANOVA was used to test the evolution of responses along training trails. Second, a two-way repeated measure ANOVA was used to compare the reinforcement type (CS+ and CS-) and the reinforcement type x trial interaction. Although ANOVA is usually not allowed in case of dichotomous data such as the MLR, Monte Carlo studies have shown that ANOVA can be used under certain conditions (Lunney, 1970), which all are met by the experiments reported here. Third, Chi^2^-tests were used for i) comparison of responses to the CS+ and CS- in a given trial, ii) comparison of spontaneous responses and retention tests, iii) comparison of the last training trial and retention tests and iv) comparison between different retention tests.

For all operant conditioning paradigms, decisions in the forced two-choice tasks were analyzed with a binomial test, since chance level of choosing one of two directions randomly was p = 0.5.

### Analysis of individuality

To analyze individuality of learning behavior we followed the analyses in Pamir et al. (2011) and Pamir et al. (2014). For the analyses in the classical conditioning paradigm we only considered animals that did not show a correct response to either the CS+ or the CS- in the very first trial and before the US was presented. Two subgroups were formed for training trials and test trial following the definition in Pamir et al. (2011). For any given trial the first subgroup included animals that expressed the correct behavior in the present trial and in the previous trial (previous correct behavior, pC). The second subgroup included animals showing the correct behavior in the present trial but did not show it in the previous trial (previous incorrect behavior, pI). The same subgroup definitions were used for the retention test with regard to a correct or incorrect response during the final training trial. We compared the two subgroups in each trial and in the retention test with a Chi^2^-test at a significance level at α = 0.05. The two subgroups are represented with upward and downward pointing triangles, respectively (Figures 5&6). Filled (open) symbols indicate that significance could (not) be established.

Following the analyses in Pamir et al. (2014), we formed separate subgroups of animals that showed a correct behavior for the first time in the second (or third) trial and tracked the subgroup behaviors across subsequent conditioning trials and the memory test. This allows to assess the robustness of the expression of a correct behavior across trials and the transfer of the behavioral expression during training to the short-term and long-term memory test situation.

Finally, in order to analyze the initiation of correct behavior, we computed for each trial the fraction of animals that responded correctly for the first time in this trial as well as the fraction of animals that never behaved correctly (non-learners, Pamir et al., 2014).

## Results

### Spontaneous response towards different odors

We first analyzed the spontaneous and naive responses to each odor (isoamyl acetate, butyric acid and cinnamaldehyde) during the very first conditioning trial before the reinforcing stimulus (US) was presented. Figure 2 shows the group averaged responses to all three odors. In all experiments approx. 60% of the animals showed a spontaneous MLR in presence of isoamyl acetate, which was significantly higher than for butyric acid and cinnamaldehyde for all experiments (Chi^2^: p < 0.001). The number of spontaneous responses to butyric acid and cinnamaldehyde was only significantly different in the first experiment (Chi^2^: p = 0.01) in which animals responded more often spontaneously to butyric acid. Isoamyl acetate is the main component of the banana blend and thus strongly associated with food and attractive for cockroaches (Lauprasert et al., 2006). This is most likely the reason for very high spontaneous response rates to isoamyl acetate. In addition we found a significant positive correlation between responses to isoamyl acetate and butyric acid in both experiments (Figure 2: 1) Phi = 0.0258, p = 0.016; 2) Phi = 0.143, p = 0.017). However, there was no significant correlation for other odor pairings which might be due to the generally low spontaneous response rates to cinnamaldehyde and butyric acid.

**Figure 2.**
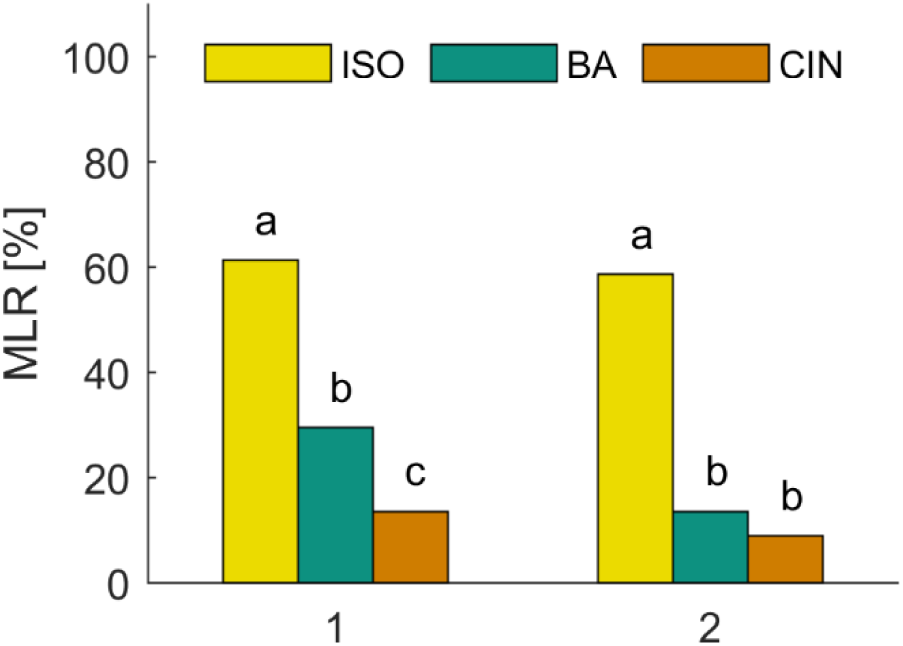
Spontaneous responses to isoamyl acetate (ISO), butyric acid (BA) and cinnamaldehyde (CIN). Analysis of the MLR in the first trial of two classical conditioning experiments show that ISO elicits significantly more spontaneous responses than BA and CIN (Chi2: p < 0.001). The spontaneous response to BA is only significantly higher compared to CIN in the first experiment (Chi2: p = 0.01).

### A novel paradigm for classical olfactory conditioning

We established a novel paradigm for classical conditioning in harnessed cockroaches (Figure 1A). The occurrence or absence of the maxilla-labia response (MLR, see Materials & Methods) was recorded as the conditioned response (CR) behavior. In this study, we used different protocols for differential olfactory conditioning (Figure 1 C&D) to investigate the expression of the CR during learning and memory retention at two different time-points.

In a first protocol we tested whether cockroaches are able to associate an odor with a reward or punishment during five consecutive training trials (inter-trial interval 10 min) followed by a retention test (after 10 min). We trained two groups of animals for which the odors isoamyl acetate and butyric acid were presented as CS+ and CS- with reversed contingencies (Figure 3A&B, respectively). The two odors did not elicit the same level of spontaneous responses (cf. section 3.1.). Due to the high initial spontaneous response to isoamyl acetate, the average level of MLR was consistently high and did not significantly increase across the five trials when isoamyl acetate was used as CS+ (Figure 3A). However, responses to the punished odor (CS-, butyric acid) decreased significantly (one-way-ANOVA: F_4, 260_ = 4.23; p = 0.002).

**Figure 3.**
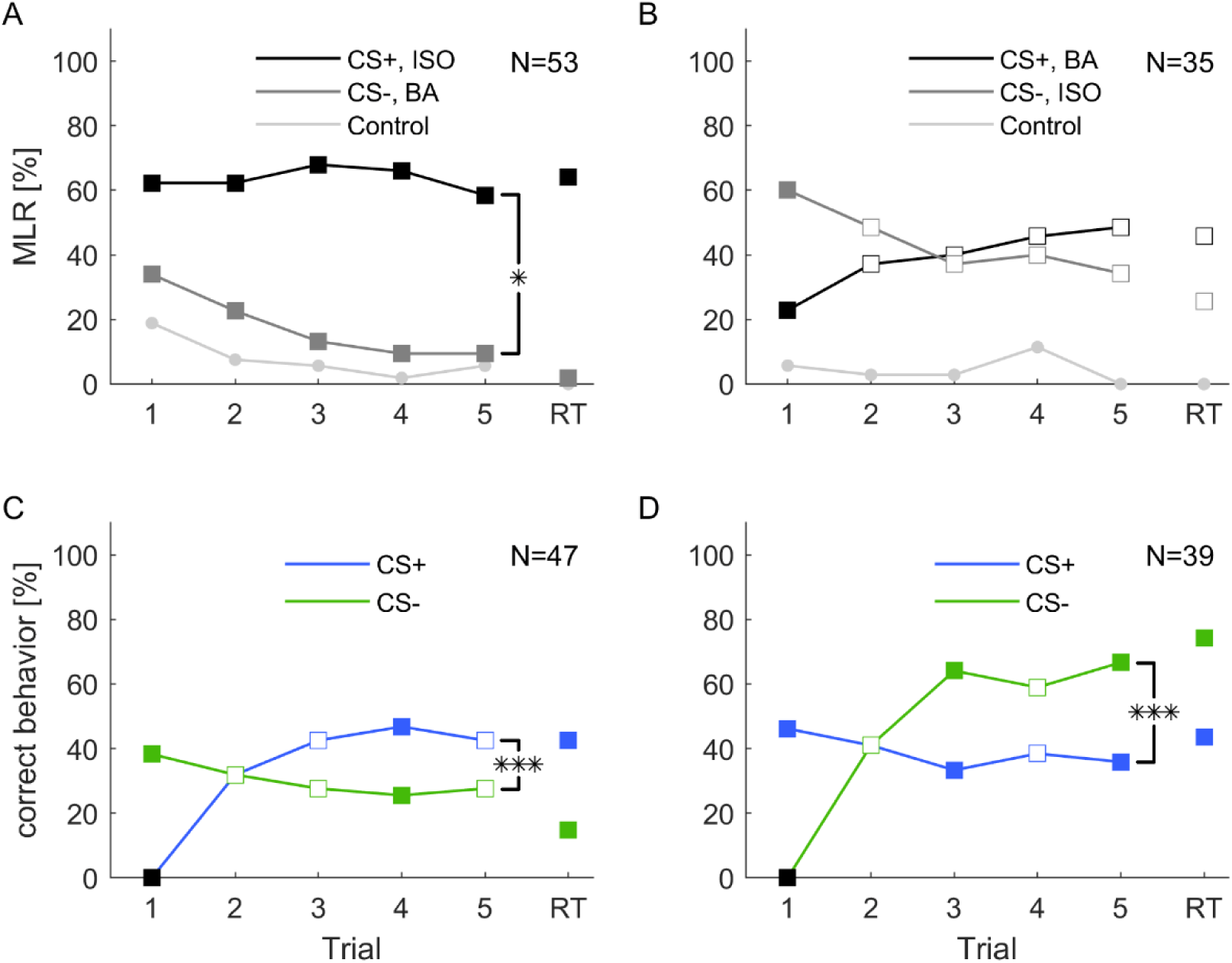
Classical olfactory conditioning over five trials with a memory retention test after 10 min. Filled squares indicate trials in which a significant difference was found between CS+ and CS- (Chi^2^ test). Asterisks indicate that responses changed for the conditioned stimuli (CS+ and CS-) over trials (2-way ANOVA, repeated measures: trea™ent x trial interaction *: p < 0.05; ***: p < 0.001). A) When isoamyl acetate (ISO) was the rewarded odor (CS+, black), the average group response did not increase during training. Responses to the punished odor BA (CS-, dark gray) decreased significantly over trials (one-way ANOVA: p = 0.002) and odor x trial interaction was significant (2-way repeated measures ANOVA: p = 0.02). B) When BA was rewarded (CS+, black), responses increased considerably, however not significantly (one-way ANOVA: p = 0.208). Responses to ISO decreased, but again, not significantly (one-way ANOVA: p = 0.189). The odor x trial interaction was not significant (2-way repeated measures ANOVA: p = 0.113). C) When excluding all spontaneous responding animals to the rewarded odor, response to CS+ increased significantly (one-way ANOVA: p < 0.001). Responses to the CS- decreased over trials, but not significantly (one-way ANOVA: p = 0.686). The trea™ent x trial interaction was significant (2-way repeated measures ANOVA: p < 0.001). D) When excluding all not spontaneous responding animals to the punished odor the number of animals that behaved correctly to CS- over trials increased significantly (one-way ANOVA: p < 0.001), while response to CS+ did not change significantly over trials (one-way ANOVA: p = 0.814). The trea™ent x trial interaction was significant (2-way repeated measures ANOVA: p < 0.001). Black squares in A&C indicate the exclusion of animals that did not behave correctly in the first trial.

When butyric acid was used as CS+, responses showed a tendency to increase over the training trials. In this case, responses to isoamyl acetate as CS-slightly decreased, however, this effect was not significant over the five trials. Animals still showed approximately 30 % responses to the CS- in the retention test.

Responses to the control odor cinnamaldehyde only decreased significantly when isoamyl acetate was the CS+ (one-way-ANOVA: F_4, 260_ = 3.11; p = 0.016), but did not change, when butyric acid was used as CS+.

Overall, responses to the CS+ and CS-differed significantly when isoamyl acetate was rewarded (2-way repeated measures ANOVA: F_4, 49_ = 3.095; p < 0.02). When butyric acid was rewarded, CS+ and CS- did not differ significantly over trials (Figure 3B).

For further analyses we pooled all animals and excluded those that did not behave correctly in the first trial respectively. In both cases correct behavior increased significantly across training trials (one-way ANOVA: CS+: F_4, 230_ = 8.808; p < 0.001; CS-: F_4, 190_ = 15.544; p < 0.001). However, neither the behavior to CS- nor the behavior to the control odor changed significantly over trials. Moreover, the interaction between trial and trea™ent was significant for CS+ (2-way repeated measures ANOVA: F_4, 43_ = 12.156, p < 0.001) and CS- (2-way repeated measures ANOVA: F_4, 35_ = 17.591, p < 0.001) and in both cases the behavior in retention tests were significantly different from each other (Figure 3 C&D).

In addition, we excluded animals that did not behave correctly respectively to CS+, CS- and the control odor. Accordingly, we could see that the effect of increasing correct behavior over trials was not only due to the exclusion of spontaneous responding or not responding animals (see Supplementary Figure 2).

### Expression of short-term and long-term memory after classical conditioning

To test memory retention after differential classical conditioning at a short- and long-term range we conducted a new experiment. A group of cockroaches were differentially trained during three consecutive trials. The group was then split in half and retention tests were performed either 1 h after the last training trial or 24 h after. In Figure 4 we show the training trials as unseparated groups, but the statistical analysis that include the training trials were conducted with splitted groups, that are shown in Supplementary Figure 1. The experiment was repeated with reversed contingencies of the odors. Responses to mechanical air stimulation (filter paper alone) did not vary and always stayed below 1.5 %.

**Figure 4.**
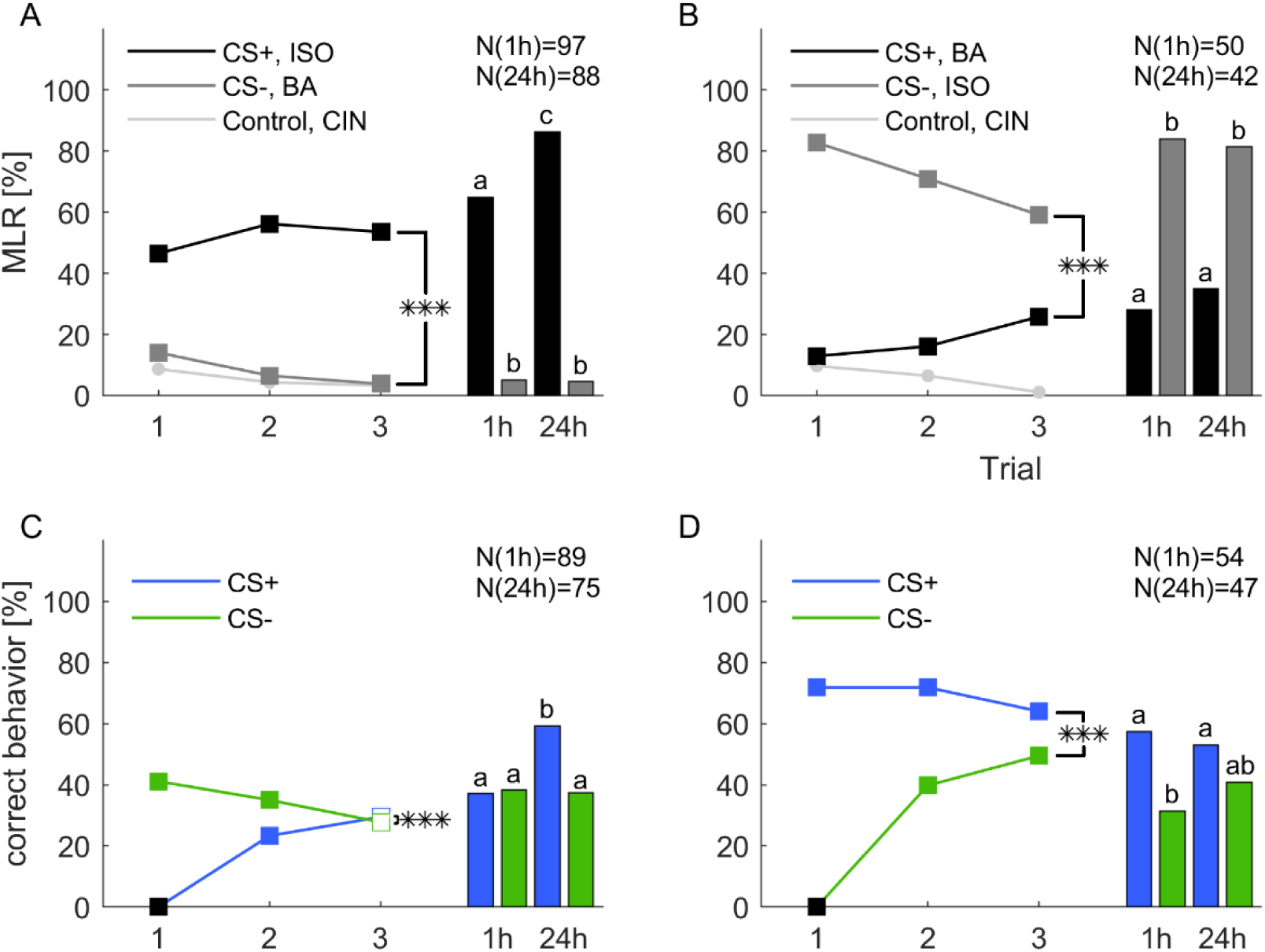
Classical olfactory conditioning over three trials with memory retention tests after 1 h and 24 h. Filled squares indicate trials in which a significant difference was found between CS+ and CS- (Chi^2^ test). Asterisks indicate a significant trea™ent x trial interaction (***: p < 0.001). Letters above the bars indicate significant differences between retention tests (Chi^2^). A) When isoamyl acetate (ISO) was rewarded (CS+, black) and butyric acid (BA) punished (CS-, dark gray), responses significantly decreased for the CS- (one-way ANOVA: p < 0.001), but did not change for the CS+ (one-way ANOVA: p = 0.155). Overall, the odor x trial interaction was significant (2-way repeated measures ANOVA: p < 0.001). B) When butyric acid was rewarded (CS+, black) and isoamyl acetate was punished (CS-, dark gray), responses to the CS+ increased, however, not significantly (one-way ANOVA: p = 0.06), but decreased significantly for the CS- (one-way ANOVA: p = 0.002). The odor x trial interaction was again significant (2-way repeated measures ANOVA: p < 0.001). C) When excluding all spontaneous responding animals to the rewarded odor the response to CS+ increased significantly (one-way ANOVA: p < 0.001). The response to CS- decreased significantly over trials (one-way ANOVA: p = 0.029). The trea™ent x trial interaction was significant (2-way repeated measures ANOVA: p < 0.001). D) When excluding all not spontaneous responding animals to the punished odor the number of animals that behaved correctly to CS- over trials increased significantly (one-way ANOVA: p < 0.001). The response to CS+ did not change significantly over trials (one-way ANOVA: p = 0.38). The trea™ent x trial interaction was significant (2-way repeated measures ANOVA: p < 0.001). Black square in C and D indicate the exclusion of animals that did not behave correctly in the first trial.

**Figure 5.**
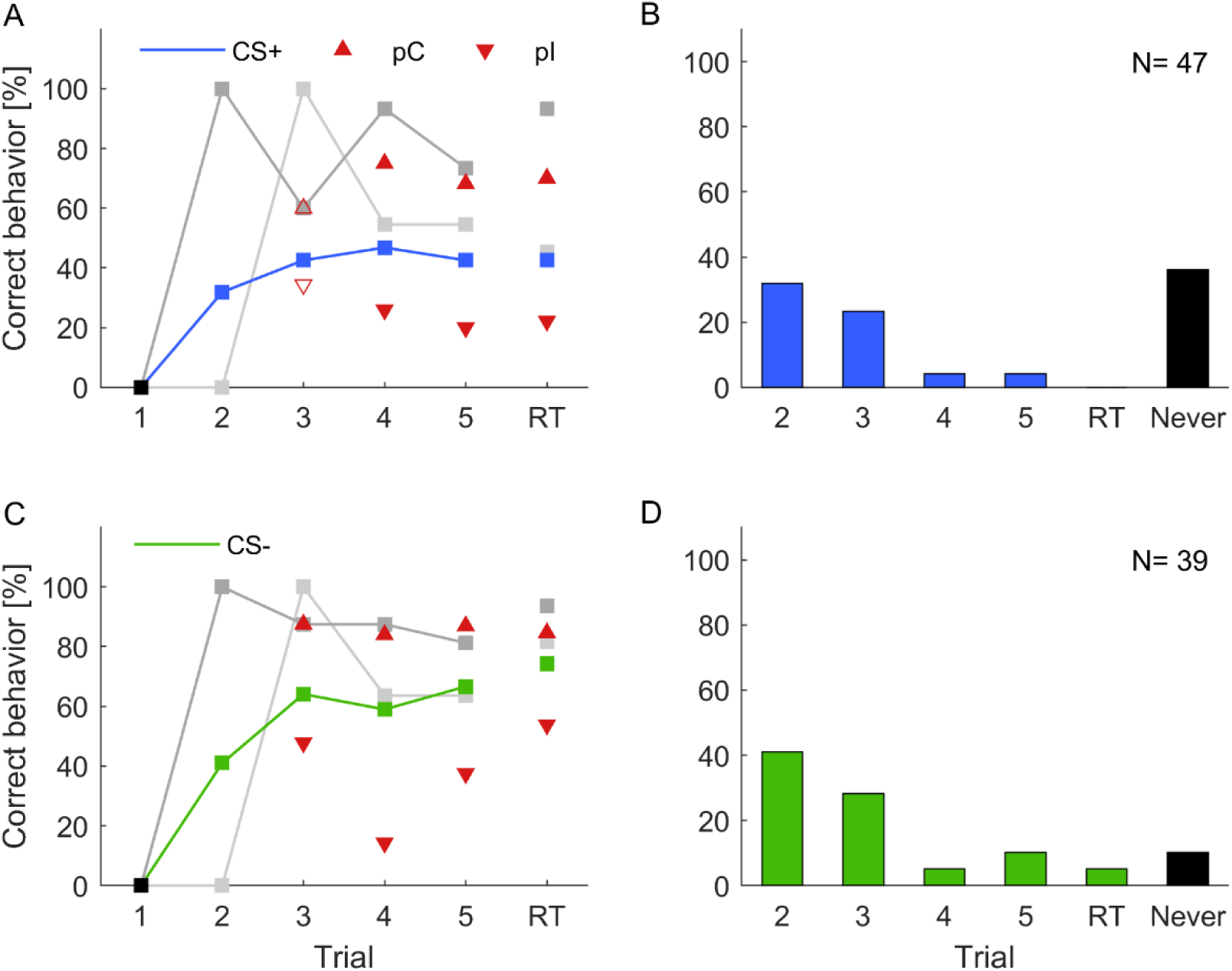
Individual learning abilities in classical olfactory conditioning. Data for isoamyl acetate as CS+ and butyric acid as CS+ were pooled, likewise data for isoamyl acetate as CS- and butyric acid as CS- were pooled. Correct learning is represented by (A) increasing responses to the rewarded odor in the case of CS+ and (C) increasing response-suppression to the punished odor in the case of CS-. (A,C) Black squares in A and C indicate the exclusion of animals that did not behave correctly in the first trial. Upward pointing triangles depict animals, which correctly behaved in the respective trial and already showed a correct behavior in the previous trial (pC). Triangles pointing downward depict animals that correctly behaved in the respective trial but did not do so in the previous trial (pI). Filled triangles indicate a significant difference of subgroups in the respective trials, empty triangles represent subgroups that were not significantly different (Chi^2^; α= 0.05). Animals behaving correctly in trial two, i.e. after a single training trial (dark gray; CS+: N = 15; CS-: N = 16) showed consistently high probabilities of correct responses in subsequent trials (cf. Pamir et al., 2014). Similarly, high response scores were observed for animals responding correctly for the first time in trial three (light gray; CS+: N = 11; CS-: N = 11). (B,D) Proportion of animals behaving correctly for the first time in the indicated trial. A high percentage of animals expressed a correct behavior already after a single training trial. A substantial proportion of animals never behaved correctly (black bars).

Overall, responses to the CS+ were significantly different from the CS-across three training trials in both groups (2-way repeated measures ANOVA: Figure 4A: F_2, 183_ = 9.266, p < 0.001; Figure 4B: F_2, 91_ = 13.016; p < 0.001). Responses to the CS-decreased significantly in both experiments (one-way ANOVA: Figure 4A: F_2, 552_ = 7.181; p < 0.001; Figure 4B: F_2, 276_ = 3.291; p = 0.002) while responses to the CS+ did not increase significantly. Responses to the control odor cinnamaldehyde decreased significantly only when butyric acid was used as CS+ (one-way ANOVA: F_2, 276_ = 3.291; p = 0.039).

During the 1 h test animals that received isoamyl acetate as CS+ maintained the elevated response level as group averaged performance, thus the retention test after 1 h was not significantly different to the response level in the last training trial (Figure 4A, Chi^2^: p = 0.143). Interestingly, the response level to isoamyl acetate was significantly higher in the 24 h retention test compared to the 1 h retention test (Chi^2^: p < 0.001), as well as in comparison to the response level at the end of training (Chi^2^: p< 0.001). When butyric acid was the rewarded odor (Figure 4B), response levels to the CS+ in both memory tests (after 1 h and 24 h) were not different from each other (Chi^2^: p = 0.475), nor from the response level at the end of training (Chi^2^: 1h: p = 0.826; 24 h: p = 0.149). The response level to the CS- resumed the initial high spontaneous response levels to isoamyl acetate during 1 h and 24 h retention. All results are summarized in Supplementary Table 2.

For the next step of analysis, animals that did not behave correctly in the first trial were excluded and for both CS+ and CS- the percentage of correct behaving animals increased (Figure 4 C&D, one-way ANOVA: CS+: F_2, 537_ = 33.537; p < 0.001; CS-: F_2, 306_ = 43.027; p < 0.001). The only other significant effect was the decrease of correct behavior to the CS- when all correct responding animals were excluded (one-way ANOVA: F_2, 537_ = 3.569; p = 0.029). Moreover, the interaction between trial and trea™ent was significant in both cases (2-way repeated measures ANOVA: Figure 4 C: F_2, 178_ = 46.719; p < 0.001; Figure 4 D: F_2, 101_ = 39.158; p < 0.001).

When excluding all spontaneously responding animals, performance in the retention test stayed at the same level as at the end of training. However, the increase of the CS+ retention test after 24 h was significant (Chi^2^: p < 0.001). After exclusion of the nonspontaneous responders, performance in all retention tests stayed similar compared to the third trial of training. However, the percentage of correct behaving animals to the CS- decreased after 1h (Chi^2^: p < 0.001) and 24 h (Chi^2^: p < 0.001).

### Individual learning performance during classical conditioning

To test whether differences in learning performance exist among individual animals we followed the analyses suggested in Pamir et al. (2011, 2014). Each of the two groups trained in the five trial classical conditioning experiment (Figure 3C&D) were divided into two subgroups (cf. Materials and Methods): a) individuals that behaved correctly in two consecutive trials (previous correct behavior, pC), and b) individuals that did not behave correctly in the previous trial but started behaving correctly in the present trial (previous incorrect behavior, pI). Previous correct behaving animals always showed a higher level of correct behavior than the average correct behavior across all animals (Figure 5, red upward triangles up), while the previous incorrect behaving individuals always showed lower correct behavior (Figure 5, red downward triangles).

Next, we analyzed the across-trial behavior of those animals that showed their first correct behavior already in the second trial (i.e. after only a single pairing of CS and US) and find that this subgroup showed consistently high rates of correct response across all trials and during retention with retention scores above 90%, both for CS+ and CS- (dark gray curves in Fig. 5A&C). Individuals that started to respond correctly in the third trial (after two pairings of CS and US, light gray curves in Fig. 5A&C) showed lower correct response levels than those animals that had started in the first trial but, still, these were comparably high considering the fact that the average response levels (blue and green curve in Figure 5A and B, respectively) included also the high performance group (dark gray curves). This indicates that fast learners are also good learners and parallels previous findings in the honeybee (Pamir et al., 2014).

In the histograms of Figure 5B&D we counted for each trial separately, how many animals responded correctly for the first time in that trial. Most animals behaved correctly for the first time after a single conditioning trial. The second largest group behaved correctly for the first time after two conditioning trials. However, a substantial portion of animals never behaved correctly (black bars in Figure 5B&D) and this group is larger for a correct behavior towards the CS+.

### Individuality in operant learning

We then tested learning, memory retention and individual differences in an operant conditioning task. Cockroaches were trained to avoid a punishment and were tested for their memory for up to 24 h. For this, we designed a forced two-choice paradigm where an individual cockroach is placed in a T-maze during repeated training trials (Figure 1B, cf. Materials & Methods). In each trial, the cockroach was allowed to choose one of the arms and entered a target box. In the first trial and irrespective of the animal’s choice, it experienced an aversive bright light stimulus. Whenever the animal chose this same side in subsequent trials, the same aversive stimulus was elicited. Learning was thus expressed in avoiding the side (left or right) that resulted in the punishment with the bright light stimulus.

In a first experiment, animals were trained for five consecutive trials and short-term memory retention was tested 35 minutes after the last trial. In a second experiment, animals were trained for three trials and a long-term retention test was performed 24 h later (Figure 1E). Animals in the first group significantly learned to avoid the punished side from the third trial on (binomial tests: p < 0.01). Animals showed correct memory for the punished side in the retention test 35 minutes after (Figure 6A). Cockroaches in the second group did not significantly show learning after two training trials. However, memory for the correct side was expressed in the 24 h retention test (binomial test: p = 0.014, Figure 6B).

**Figure 6.**
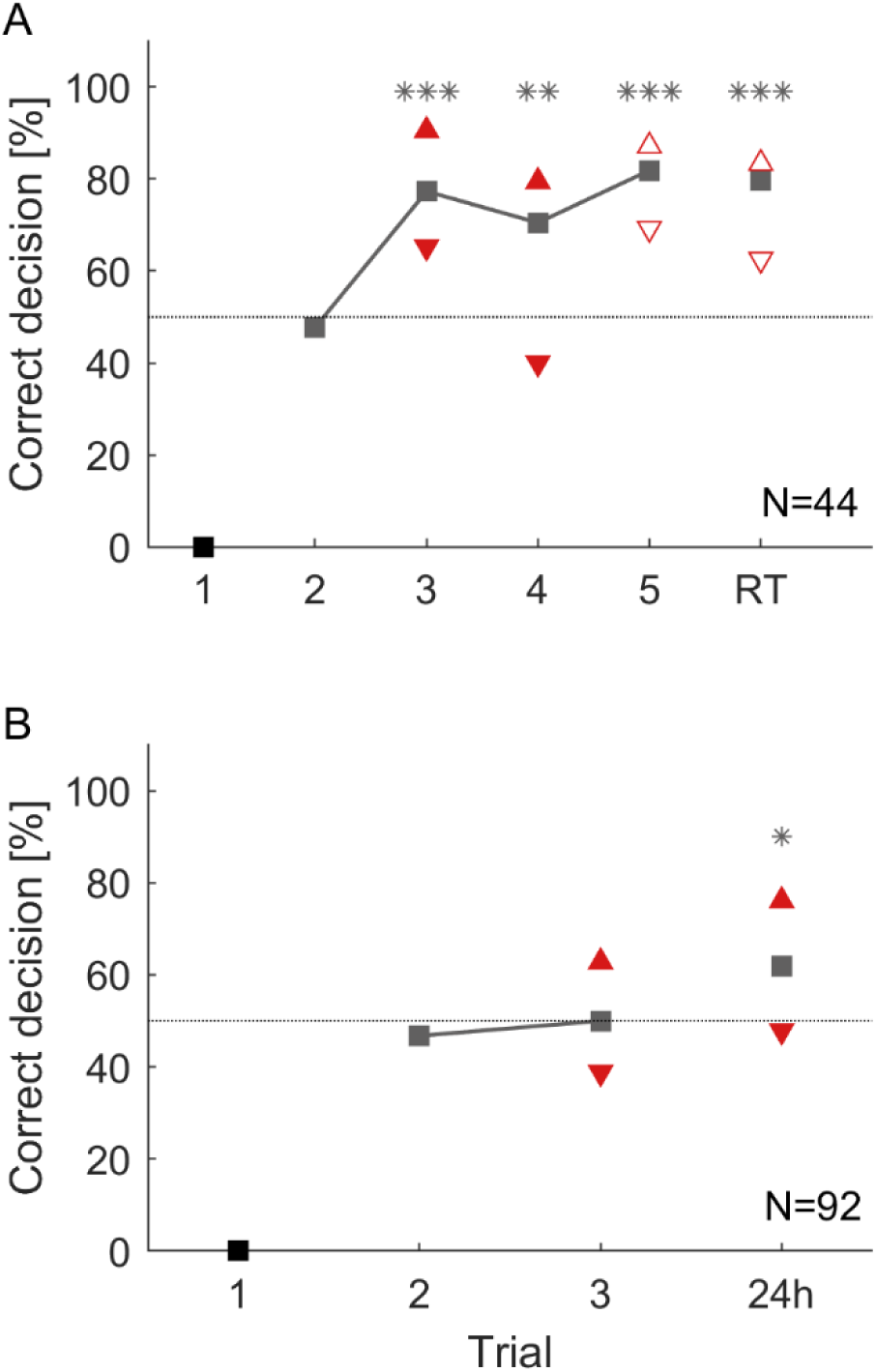
Overall group performance and individual choices during an operant conditioning task in a T-maze A) over five consecutive trials and a 35 min retention test (RT). From the third trial on cockroaches chose significantly more often the correct direction (**: p < 0.01; ***: p < 0.001). B) Operant conditioning over three trials and a retention test after 24 h. However, this time cockroaches did not learn during training but chose significantly more the correct direction in the 24 h retention test (*: p < 0.05). Triangles that point upwards depict animals that chose the correct direction in two consecutive trials (pC). Triangles that show downwards depict animals that chose the correct direction in the present trial but did not choose it in the previous trial (pI). Filled triangles differ significantly from each other within a trial and empty ones do not (Chi^2^; α = 0.05).

To study individual differences in this operant learning and memory tasks, animals were again attributed to two subgroups. In the short-term memory experiment, animals in the subgroup showing the correct behavior in two consecutive trials (pC) always performed better than the group average while animals in the subgroup pI consistently showed fewer correct choices in the present trial. Behavioral choices of previous correct deciding animals (pC) and previous incorrect deciding animals (pI) significantly differed in the third and fourth trial during training (Chi^2^: p < 0.05). This difference was not significant in the fifth learning trial, nor in the retention test after 35 min (Chi^2^: trial 3: p = 0.161; trial 4: p = 0.186, Figure 6A).

In the long-term memory experiment, the two subgroups (pC and pI) again differed significantly after two training trials. During the 24 h retention test the subgroup of animals that had shown a correct decision in the last training trial significantly outperformed those animals that had made an incorrect decision in the last training trial (Figure 6B: Chi^2^: p < 0.05).

## Discussion

In the present work we show that the adult American cockroach, *Periplaneta americana* can solve classical olfactory and operant spatial conditioning tasks. In both cases, animals could learn to establish a conditioned response to the rewarded stimulus (CS+), and to diminish their responses to the punished odor (CS-) despite the fact that, in the classical conditioning task, the two odors were not equally important to the animals (high spontaneous responses to isoamyl acetate). Overall, training resulted in the successful expression of short term memory and long term memory (after 24h) in both conditioning tasks. We further show that cockroaches express individuality in their learning and memory performance in classical and operant conditioning.

### Classical olfactory conditioning in the cockroach

For the present study we established a novel classical conditioning paradigm in harnessed cockroaches that allows to observe the expression (or non-expression) of a discrete conditioned response behavior, the <u>m</u>axilla labia response (MLR) during learning and memory retention. The development of this paradigm was inspired by the highly successful proboscis extension reflex (PER) paradigm in the honeybee (e.g. (Bitterman et al., 1983; Giurfa and Sandoz, 2012; Kuwabara, 1957; Menzel, 2012; Takeda, 1961) in which the extension (or non-extension) of the proboscis is observed as a discrete conditioned response behavior.

A number of previous studies have investigated classical conditioning in the cockroach using training and test conditions that differ fundamentally from our MLR paradigm. In studies by Watanabe et al. (2003), Sato et al. (2006) and Liu and Sakuma (2013), in the German cockroach, unrestrained cockroaches were placed in a cylindrical chamber during repeated conditioning trials with one odor paired with sucrose reward (CS+) and a second odor paired with salt punishment (CS-). Sato et al. (2006) could prove that beyond simple olfactory discrimination learning, cockroaches exhibited excellent learning performance in an occasion setting paradigm in which a visual context defines the contingency between olfactory CSs (conditioning stimuli) and gustatory USs (unconditioned stimuli). Watanabe et al. (2003) extended their classical conditioning protocol in unrestrained cockroaches to harnessed cockroaches that were subsequently tested under freely moving conditions in a test arena where they could choose between the two previously conditioned odors. This paradigm, however, did not establish a clear conditioned response observable during training and thus expression of a conditioned response behavior is only accessible during memory retention and under conditions different from training. Classical conditioning leads to an increase in response of salivary neurons to an odor associated with sucrose reward in the cockroach (Watanabe and Mizunami, 2006). After differential conditioning one odor paired with sucrose and another odor without reward, the sucrose-associated odor induced an increase in the level of salivation, but the odor presented alone did not, proving classical conditioning of salivation in cockroaches (Watanabe and Mizunami, 2007). Classical conditioning of salivation has first been shown a century ago by Pavlov in his famous dog experiments (Pavlov, 1927). Restrained cockroaches were further used to study spatial (e.g. Kwon et al., 2004) or visual-olfactory associative learning and memory (e.g. Lent et al., 2007; Lent and Kwon, 2004; Pintér et al., 2005) by quantifying the antennal projection response (APR) of animals that were tethered in the middle of an arena (Pomaville and Lent, 2018). The APR is based on the observation that antennal motor actions can be elicited by different modalities, including olfactory, tactile and visual stimuli (e.g. Erber et al., 1997; Menzel et al., 1994). Conditioning the APR consists in quantifying directed antennal movements towards the direction of a rewarded visual stimulus and was inspired by operant conditioning of bees to extend their antennae towards a target in order to receive a reward (e.g. Kisch and Erber, 1999; Menzel et al., 1994). The advantage of training immobilized insects provides a powerful technique for studying the neuronal basis (by e.g. employing neurophysiological and pharmacological techniques) of learning and memory in a simpler nervous system compared to vertebrates.

### Initial response behavior during classical conditioning

Stimuli used for studying olfactory learning and memory in insects mostly employ odors which are relevant in the natural context, such as communication signals (i.e. pheromones) or food-related odors. Isoamyl acetate constitutes the most salient compound of the banana blend and is perceived as the smell of banana (Schubert et al., 2014). This odor is clearly food related and thus highly attractive for cockroaches (Lauprasert et al., 2006). This likely explains why, in our olfactory conditioning experiments, we observed a high level of initial responses to isoamyl acetate in the first trial (Figure 2). Consequently it was difficult to observe learning (i.e. increasing conditioned response levels) when this odor was paired with a sucrose reward (CS+) since response levels were consistently high (> ∼60%) from the first trial on and throughout training (Figure 3A). In the 24 h retention test, however, the MLR to isoamyl acetate was significantly increased compared to the response in the last training trial (Figure 4A), indicating that a long-term memory had been established. When isoamyl acetate was paired with salt punishment (CS-) animals learned to suppress their responses during training (Figure 4B). Initial responses to butyric acid were significantly lower (∼ 30%) at the beginning of training in all cases (Figure 2). When associated to sugar, responses increased but never exceeded 50% even after five training trials. Spontaneous responses to butyric acid were completely abolished during training and memory retention when paired with punishment (Figure 3B). Concluding, the two odors employed in our study were not equally attractive to the animals.

### Operant spatial conditioning in cockroaches

A frequently used setup for operant conditioning is a Y- or T-maze, which is extensively used to study operant learning and decision making in rodents. T- or Y-maze (dual choice) experiments in invertebrates have been used broadly to study visual or olfactory absolute and differential learning in free flying bees (e.g. for review: Avarguès-Weber et al., 2011; Giurfa et al., 1999, 2001; Nouvian and Galizia, 2019; Srinivasan et al., 1998), in ants (e.g. Camlitepe and Aksoy, 2010; Dupuy et al., 2006) and in wasps (Hoedjes et al., 2012). In cockroaches, operant learning has repeatedly been studied in open arenas (e.g. Balderrama, 1980; Sakura et al., 2002; Sakura and Mizunami, 2001). The first work on operant conditioning in cockroaches was carried out by Balderrama (1980) who trained free moving cockroaches individually in a simple training chamber to associate two artificial odors to sucrose and salt solutions, respectively, and testing discriminatory learning performance by measuring the odor preference before and after training. Spontaneous preference for one of the odors before training could be modified already with one trial and retention lasted up to 7 days. To date, there are only two studies that have challenged cockroaches in T-maze tasks, the first testing the influence of feces pheromones on directional orientation (Bell et al., 1973), while the second investigated effects of protein synthesis inhibiting drugs on learning and retention by training animals to avoid shock on one of the sides (Barraco et al., 1981). Our reason to perform an operant learning paradigm in the T-maze was to establish a forced binary choice that can be analyzed during acquisition and memory retention in a defined trial design. Electric shock as used for a punishing stimulus in the previous study by Barraco et al. (1981) seems a rather unnatural aversive stimulus that is unlikely to appear in nature. We decided to use bright light as negative reinforcer since cockroaches naturally avoid bright light and seek shelter in darkness (Turner, 1912).

Cockroaches started to avoid the side that was punished after a few trials. However, training results were variable across the two experiments. Previous studies concluded that cockroaches show unpredicted searching behavior (Balderrama, 1980). Similarly, we could observe different traits in behavior, which might partly underlie the variance in choice behavior. For example, some cockroaches show a high explorative behavior, possibly in search for an exit from the maze, and these did not seem to care much about the reinforcing stimulus while others stayed almost immobile throughout a trial and moved little. The punishing effect of light is limited because it has no harming consequence for the animal. They may thus habituate to the aversive light stimulus. The T-maze experiments in Barraco et al. (1981) using electric shock as negative reinforcer resulted in surprisingly high correct choice rates. However, a strong light seems to be repellent for most cockroaches since they normally try to hide in a dark place when e.g. the light in a room is switched on. In future experiments we want to explore whether a paradigm for appetitive operant conditioning can lead to higher levels of correct choice performance in cockroaches.

### Individual behavioral expression of learning and memory

Our approach to study individuality in learning performance during classical conditioning was inspired by two previous studies by Pamir et al. (2011, 2014) that investigated a large number of data sets on classical appetitive conditioning in the honeybee. The authors were able to extract from the immense amount of data that honeybees express individual learning behavior and that a group of animals can be separated into at least two subgroups, learners and non-learners. Both studies by Pamir and colleagues have investigated behavioral learning and memory expression only towards the CS+. We have extended their analysis including behavioral learning and memory expression towards the CS- (Figure 5C&D).

As for the honeybee (Pamir et al., 2014), a large fraction of animals (>35%) never showed the correct behavior to the CS+ odor in any of the learning trials or the retention test (Figure 5B&D). These animals may be considered non-learners. When taking into account only those animals that expressed the correct conditioned behavior at least once during the training session, then we find that those animals expressed this behavior for the first time after average 1.7 conditioning trials towards the CS+ and after average 1.8 conditioning trials towards the CS-. Indeed, the largest fraction (50%) of responding animals showed a correct conditioned response behavior for the first time already after a single conditioning trial (single-trial learning), both towards the CS+ and the CS-. In effect, 86,6 % of learners showed a first correct behavior to the CS+ or CS- already after the first or second conditioning trial, indicating rapid learning after a single or two trials. These numbers match closely those obtained in the honeybee where typically ∼50% of individuals in a group of honeybees showed a conditioned response after a single training trial (Pamir et al., 2014). Moreover, the correct expression of learned behavior in fast learners is remarkably stable as can be seen when following the across-trial CR behavior of the subgroup of cockroaches that showed a correct behavior after a single conditioning trial (dark gray curve in Figure 5A&C). When looking at short-term memory retention in those animals (Figure 5A&C), 93.3% and 93.8% expressed the correct behavior during the test for CS+ and CS-, respectively. Conversely, of the fraction of animals that showed the correct behavior during short-term memory retention for CS+ and CS-, 95% and 82.8%, respectively, were fast learning individuals expressing the correct behavior after a single or two training trials. Similarly, Pamir et al. (2014) reported that honeybees that responded earlier showed a higher long-term memory retention than those responding later. Taken together our results indicate that (i) individual cockroaches are able to learn efficiently during only one or two conditioning trials, and (ii) fast learners are also good learners that robustly express the correct behavior throughout the training session and achieve very high retention scores.

Thus, in line with the results on honeybees reported by Pamir et al (2011, 2014), we conclude from our results that the gradually increasing group-average learning curve does not adequately represent the behavior of individual animals. Rather, it confounds three attributes of individual learning: the ability or inability to learn a given task (learners vs. non-learners), the fast acquisition of a correct conditioned response behavior in learners, and a high robustness of the conditioned response expression during consecutive training and memory retention trials. Moreover, we could establish the same general result in an operant learning task in the cockroach. The latter result is in line with a study in bumblebees (Muller and Chittka, 2012) observing that some individuals were consistently better than others in associating different cues with reward or punishment in an operant learning task.

Interestingly, these congruent results in the honeybee and cockroach, two evolutionary far separated species, are in contrast to the long-standing notion on learning abilities in fruit flies. An early report on olfactory learning in *Drosophila melanogaster* by Quinn et al. (1974) using a meanwhile well-established and heavily used group assay for classical olfactory conditioning of flies concluded that the expression of behavior in the individual was probabilistic such that a group of flies can be treated as homogeneous with respect to the ability to acquire a correct CR behavior. This notion has been challenged by a more recent study (Chabaud et al., 2010), but awaits further conclusive investigation. We hypothesize that fruit flies exhibit individual learning performance that are very similar to those observed in the honeybee and established for the cockroach in this study.

### Possible causes for the individual expression of learned behavior

What could be the underlying causes for the observed individuality in behavioral learning performance? At the neuronal circuit level, learning-induced plasticity has been observed at different sites within the system. Two studies in honeybees found correlations between the behavioral performance in individuals and the expression of plasticity in the nervous system. Rath et al. (2011) performed Ca-imaging in the projection neurons of the antennal lobe. For their analysis they formed two subgroups of learners and non-learners based on their conditioned response behavior and reported that, as a result of classical olfactory conditioning, odor response patterns in the projection neuron population became more distinct in learners but not in non-learners. Haenicke et al. (2018) performed Ca-imaging from the projection neuron boutons in the mushroom body calyx of the honeybee and found that the level of neuronal plasticity correlates significantly with the level of behavioral plasticity across individual animals in classical olfactory conditioning. Mushroom body output neurons have been shown to convey the valence of odors following classical conditioning in bees (Strube-Bloss et al., 2011, 2016) and flies (e.g. Aso et al., 2014; Hige et al., 2015). In bees, the level of observed plasticity in these neurons after classical conditioning again correlates with the behavioral performance during the retention test (Strube-Bloss, d’Albis, Menzel & Nawrot, unpublished data). Thus, individuality in the conditioned response performance during memory retention has been linked to the underlying plasticity in the neural circuitry.

In bees, a significant correlation between their sensitivity to sucrose concentration and learning performance during an olfactory task has been reported (Scheiner et al., 2004). Pamir et al. (2014) re-analyzed data from Scheiner et al., (2001) showing that sucrose responsiveness, interpreted as a proxy to the state of satiety, correlates with learning performance, both in olfactory and tactile classical conditioning.

In addition, a number of studies have linked variations in learning abilities with genetic variation across individuals. In the honeybee, for example, animals which performed well in olfactory/mechanosensory conditioning also performed well in visual learning. On the other hand, good and poor learners from strains selected for olfactory conditioning differed significantly in their visual learning values. Thus, genetic differences exist between different strains and such genetic variation can account for differences in learning in individuals (e.g. Brandes et al., 1988; Brandes and Menzel, 1990). Another study on honey bees considering individual differences in a latent inhibition learning task (learning that some stimuli are *not* signals of important events) also proved a genetic predisposition for learning this task (Chandra et al., 2000). A study on fruit flies that learned to associate a chemical cue (quinine) with a particular substrate showed that individuals still avoided this substrate several hours after the cue had been removed, were expected to contribute more alleles to the next generation. From about generation 15 on the experimental populations showed marked ability to avoid oviposition substrates that several hours earlier had contained the chemical cue (for review see: Dukas, 2008; Mery and Kawecki, 2002). Indeed, genetic variation might underlie individuality in behavior in general and in learning behavior specifically. However, to our knowledge genetic variation has not been studied in cockroaches in relation to behavioral traits. Unfortunately, maturation and reproduction cycles in cockroaches are rather long.

### Outlook

In the present study we investigated individual learning performance and learning speed in single learning tasks (classical olfactory conditioning or operant place learning). In future studies we will extend our analyses of individuality in two directions. First, we will investigate whether the behavioral performance of individuals is consistent across different learning paradigms, i.e. whether good and fast learners in one classical conditioning paradigm will also perform above average in different classical or operant conditioning tasks. To our knowledge there is only one invertebrate study where something comparable was published with honeybees (Honegger et al., 2019). Second, we are interested in consistency across days or weeks investigating whether a high/low performance of one individual is equally high/low during a repetition of the same or similar task at a later point in time.

## Acknowledgements

C.A. receives a PhD scholarship from the Research Training Group *Neural Circuit Analysis on the Cellular and Subcellular Level* funded through the German Research Foundation (DFG-RTG 1960, grant no. 233886668). Additional funding is received from the German Research Foundation within the Research Unit *Structure, Plasticity and Behavioral Function of the Drosophila mushroom body* (DFG-FOR 2705, grant no. 403329959 to M.N.). We thank Sandra Mastani for contributions in an early stage of this project. We thank Peter Kloppenburg for discussion, sharing cockroach facilities and use of animals. We thank the team of our mechanical workshop headed by Leo Lesson, which designed and manufactured the odor stimulation devices and the flexible cockroach maze kit.

