## Supplementary Material for "Cockroaches show individuality in learning and memory during classical and operant conditioning"

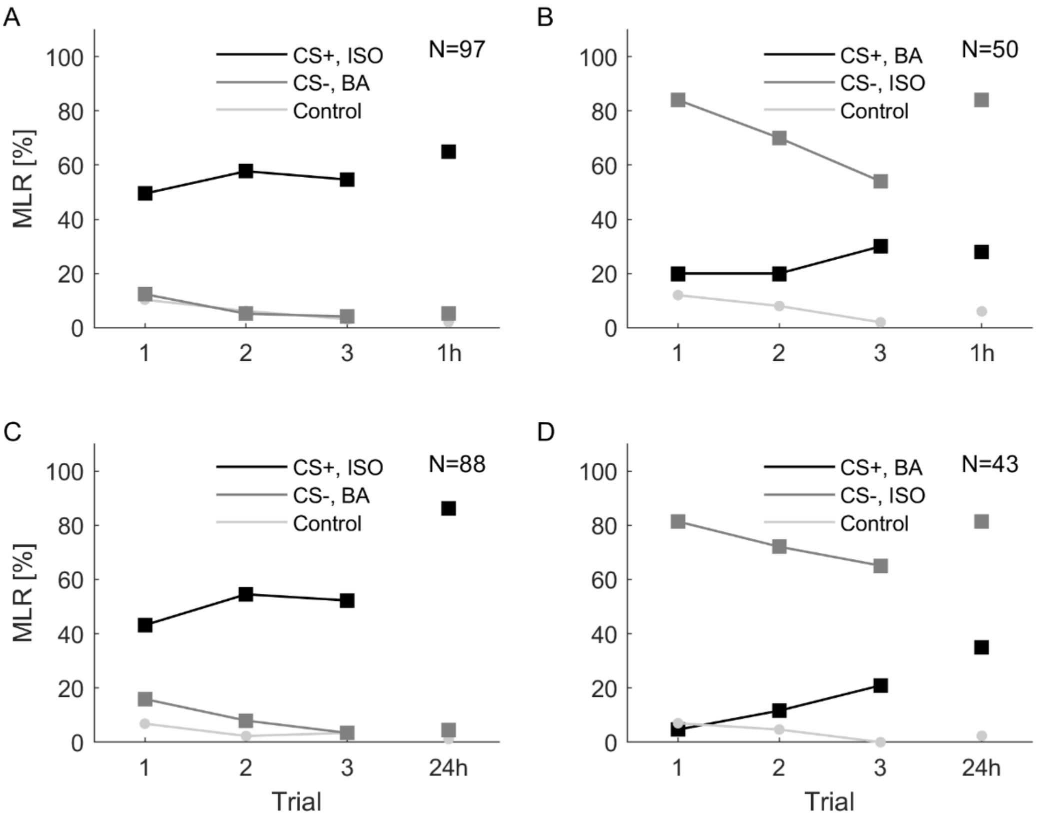


**Supplementary Figure 1** Classical olfactory conditioning over three trials with memory retention tests after 1 h and 24 h. A) Isoamyl acetate (ISO, black) was used as CS+, butyric acid (BA, dark gray) was used as CS- and cinnamaldehyde (light gray) was used as control. The retention test was after 1 h. B) BA (black) was used as CS+, ISO (dark gray) was used as CS- and cinnamaldehyde (light gray) was used as control odor. The retention test was after 1h. C) ISO (black) was used as CS+, BA (dark gray) was used as CS- and cinnamaldehyde (light gray) was used as control odor. The retention test was after 24 h. D) BA (black) was used as CS+, ISO (dark gray) was used as CS- and cinnamaldehyde (light gray) was used as control odor. The retention test was after 24 h**.**

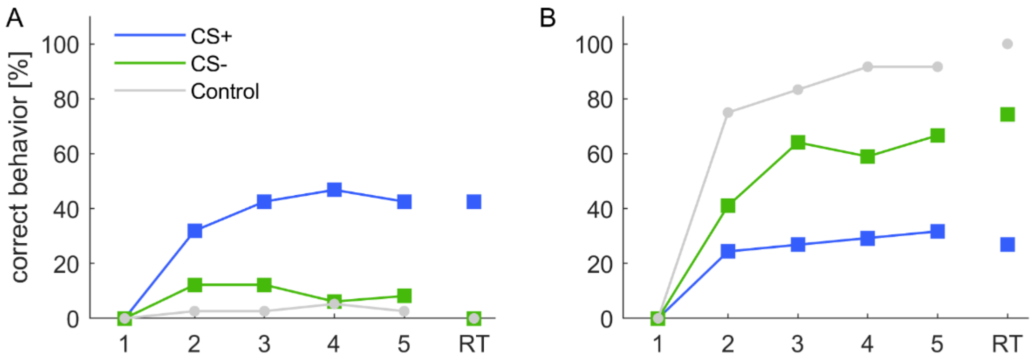


**Supplementary Figure 2** Classical olfactory conditioning over five trials with a memory retention test after 10 minutes. A) Animals that did respond spontaneously in the first trial to the respective stimulus were excluded. The correct behavior to CS+ (blue) increased significantly over trials (one-way ANOVA: $p < 0.001$, N = 47) while the behavior towards the CS- (green) and control (gray) odors did not change over trials (CS-: p = 0.133, N = 49; control: p = 0.394, N = 76). B) Animals that did not respond spontaneously in the first trial to the respective stimulus were excluded. The correct behavior to CS- increased significantly over trials (one-way ANOVA: $p < 0.001$, N = 39). Unexpectedly, the behavioral expression also increased (the expression of the MLR decreased) towards the CS+ (one-way ANOVA: p = 0.003; N = 41) albeit to a lesser extent and mostly from trial one (spontaneous response) to a rather constant level represented by a small fraction of all animals. This might be explained to some extent with spontaneously responding non-learners that expressed spontaneous responses towards the CS+ odor with a low probability or with the satiety state of the animals. The observed decrease of the MLR towards the control odor is expected, as this odor was not rewarded. In many protocols for differential conditioning the CS+ odor is rewarded and the CS- odor is not rewarded which results in a decrease of the conditioned response to the non-rewarded odor (e.g. Pamir et al., 2011). Since, the sample size is too low we could not test for an increase in correct responses to the control odor (N = 12). In our experiments this corresponds with the reduction of the MLR (increase of correct behavior) towards the control odor.

**Supplementary Table 1** Comparison between training trials and retention tests. Isoamyl acetate (ISO) was used as CS+ and butyric acid (BA) was used as CS-. Data is depicted in Supplementary Figure 1 A&C. Chi² test was used to analyze differences in the number of conditioned responses between retention tests and the respective trial 1 or 3 and p-values are listed. Numbers in bold are < 0.05 and indicate significant difference between the training trial and retention test.

| | Retentiontest 1h (ISO) | Retentiontest 1h (BA) | Retentiontest24 h (ISO) | Retentiontest 24h (BA) |
|---|---|---|---|---|
| **Trial 1** | **0.03** | 0.076 | **< 0.001** | **0.0129** |
| **Trial 3** | 0.143 | 0.733 | **<0.001** | 0.7 |

**Supplementary Table 2** Comparison between training trials and retention tests. Butyric acid (BA) was used as CS+ and isoamyl acetate (ISO) was used as CS-. Data is depicted in Supplementary Figure 1 B&D. Chi² test was used to analyze differences in the number of conditioned responses between retention tests and the respective trial 1 or 3 and p-values are listed. Numbers in bold are < 0.05 and indicate significant difference between the training trial and retention test.

| | Retentiontest 1h (BA) | Retentiontest 1h (ISO) | Retentiontest 24h (BA) | Retentiontest24 h (ISO) |
|---|---|---|---|---|
| **Trial 1** | 0.349 | 1 | **< 0.001** | 1 |
| **Trial 3** | 0.826 | **0.001** | 0.149 | 0.088 |
